# Distinct proliferative and neuronal programmes of chromatin binding and gene activation by ASCL1 are cell cycle stage-specific

**DOI:** 10.1101/2024.10.07.616995

**Authors:** William F. Beckman, Lydia M. Parkinson, Lewis Chaytor, Anna Philpott

## Abstract

ASCL1 is a potent proneural factor with paradoxical functions during development, promoting both progenitor pool expansion and neuronal differentiation. How a single factor executes and switches between these potentially opposing functions remain to be understood. Using neuroblastoma cells as a model system, we show that ASCL1 exhibits cell cycle phase-dependent chromatin binding patterns. In cycling cells, S/G2/M phase-enriched binding occurs at promoters of transcribed pro-mitotic genes, while G1 phase-enriched binding of ASCL1 is associated with the priming of pro-neuronal enhancer loci. Prolonged G1 arrest is further required to activate these ASCL1-bound and primed neuronal enhancers to drive neuronal differentiation. Thus, we reveal that the same transcription factor can control distinct transcriptional programmes at different cell cycle stages, and demonstrate how lengthening of G1 allows engagement of a differentiation programme by turning unproductive factor binding into productive interactions.

## Introduction

Achaete-Scute homolog 1 (ASCL1) is a potent driver of neurogenesis in both the central and peripheral nervous system (CNS/PNS) and has emerged as a key player in the balance between neuronal progenitor cell differentiation and self-renewal. Strict spatiotemporal orchestration of these processes is essential for the development of the mammalian nervous system to ensure that a sufficient number of neurons are generated at appropriate stages, without progenitor pool depletion ^1^. ASCL1 is necessary and sufficient to activate neuronal differentiation and specify subtype identity ^2–7^. Perturbation of the physiological activity of ASCL1 results in atrophy or delayed development of CNS and PNS components ^8,9^. In the PNS, ASCL1 is transiently expressed in the precursors of enteric and sympathetic neurons, including noradrenergic neuroblasts, and is subsequently downregulated concomitantly with neuronal differentiation ^10–13^. In the CNS, ASCL1 controls fate specification of neuronal and glial lineages, where knockout impairs the normal development of multiple brain regions ^14–17^. Here, ASCL1 is also transiently expressed in mitotic neuronal precursors, with the onset of this expression coinciding with the transition of quiescent stem cells to immature progenitors ^18^, and with subsequent downregulation of ASCL1 coinciding with terminal differentiation and cell cycle exit ^14^. *Ascl1*-null mice show a reduction in the number of cycling intermediate neural progenitor cells in the subventricular zone of the ventral telencephalon, stemming from reduced expression of *Ascl1* targets encoding components of the E2F1 and FoxM1 pathways ^15^. Ascl1 was also shown to bind and activate many genes involved in cell cycle progression, such as *E2f1*, *Cdk1* and *Cdc25b* ^15^.

Neuronal differentiation is tightly and inversely linked to cell cycle progression, to the extent that many of the archetypal cell cycle regulators also have non-canonical roles in neuronal specification ^19,20^. Repurposing of these cell cycle regulators alters cell cycle structure; indeed, lengthening of G1 phase has repeatedly been shown to promote differentiation in somatic stem cells ^21–24^, and rapid transition through G1 phase is a hallmark of the pluripotent state ^25^. Mechanistically, the effect of G1 length on specification derives from the capacity of lineage-defining pioneer factors to prime cell type specific enhancers in G1 ^26^, but requires a lengthening of G1 phase to fully commission them ^27^. This primed enhancer state is associated with low levels of H3K4me1 and chromatin accessibility ^27^ where activation leads to deposition of H3K4me1, H3K27ac and increased accessibility, in part due to recruitment of SWI/SNF by eRNAs which in turn recruits MLL3/4, p300/CBP and Mediator ^28^, initiating determinate cell type trajectories.

We have previously shown that ASCL1 is capable of performing both context-dependent proliferative and differentiation functions in neuroblastoma, a paediatric cancer originating from immature sympathoadrenal precursors of the PNS ^29,30^, where ASCL1 is normally a member of the adrenergic subtype core regulatory gene circuit driving growth ^31–33^. In neuroblastoma cells, knockout of ASCL1 reduces *in vitro* proliferation rate ^12^, while overexpression drives potent cell cycle exit and neuronal differentiation ^34,35^. This ability to drive differentiation is further enhanced when ASCL1 multi-site phosphorylation is prevented ^34–36^, showing that post-translational modifications of ASCL1 in response to cell cycle-dependent CDK2 may regulate its activity, and indicating the possibility that ASCL1 function may be differentially controlled at different stages of the cell cycle. Thus, neuroblastoma cells provide a convenient model system to test whether the potentially conflicting functions of ASCL1 in proliferation and differentiation could be explained by differential activities of this transcription factor at different phases of the cell cycle. To investigate potential cell cycle stage-dependent functions of ASCl1, we have performed ChIP-seq and RNA-seq to detect ASCL1 binding and gene activation in neuroblastoma cells at different cell cycle stages. In SG2M, ASCL1 binding sites are enriched at promoters of cell cycle-associated genes, and supports their expression. In contrast, ASCL1 binding sites enriched in G1 show more association with enhancers of neuronal function genes. However, in cycling cells these G1 ASCL1-bound genes are poorly expressed and remain relatively inaccessible, showing gene target activation only after cells are arrested in G1.

## Results

### ASCL1 binds to both pro-proliferative and pro-neuronal loci in asynchronous cycling cells

We have previously shown that ASCL1 overexpression in neuroblastoma cell lines results in its widespread binding near loci associated with neuronal structures and functions, and results in subsequent neuronal differentiation. However, ASCL1 is expressed at readily detectable levels in many rapidly cycling adrenergic-type neuroblastoma cell lines where it supports their proliferation^12^. We currently have limited understanding of endogenous ASCL1 binding and how it supports gene expression in proliferating neuroblastoma cells ^37^. To investigate this, we performed ChIP-seq for ASCL1 on cycling SK-N-BE(2)-C cells, a representative MYCN-amplified neuroblastoma cell line. Approximately 100,000 high confidence ASCL1 binding sites were found to be present in 2 out of 3 replicates (Figure 1A, B). Consistent with the known role of ASCL1 in binding distal gene regulatory elements ^2^, we found that only approximately 25% of these high confidence binding sites were within 3kb of transcription start sites, with most binding sites being intergenic (Figure 1C). To identify potential direct regulatory targets of ASCL1, we limited gene annotation to the nearest expressed gene within 50kb of each peak (Figure S1A), leaving 50,488 peaks annotated to 11,950 genes. Gene ontology analysis of these bound genes in asynchronous cycling cells at all cell cycle stages revealed that ASCL1 binding is associated most prominently with core processes such as ribonucleoprotein complex biogenesis and RNA splicing, proliferative processes such as chromosome segregation and DNA replication, and pro-neuronal processes such as the regulation of neuron projection development (Figure 1D). We performed ASCL1 ChIP-seq in another MYCN-amplified neuroblastoma cell line (IMR-32) and a non MYCN-amplified cell line (SH-SY5Y) showing that ASCL1 binding at loci associated with both proliferative and neuronal genes is a general feature of ASCL1 functionality in neuroblastoma (Figure S1B).

**Figure 1.**
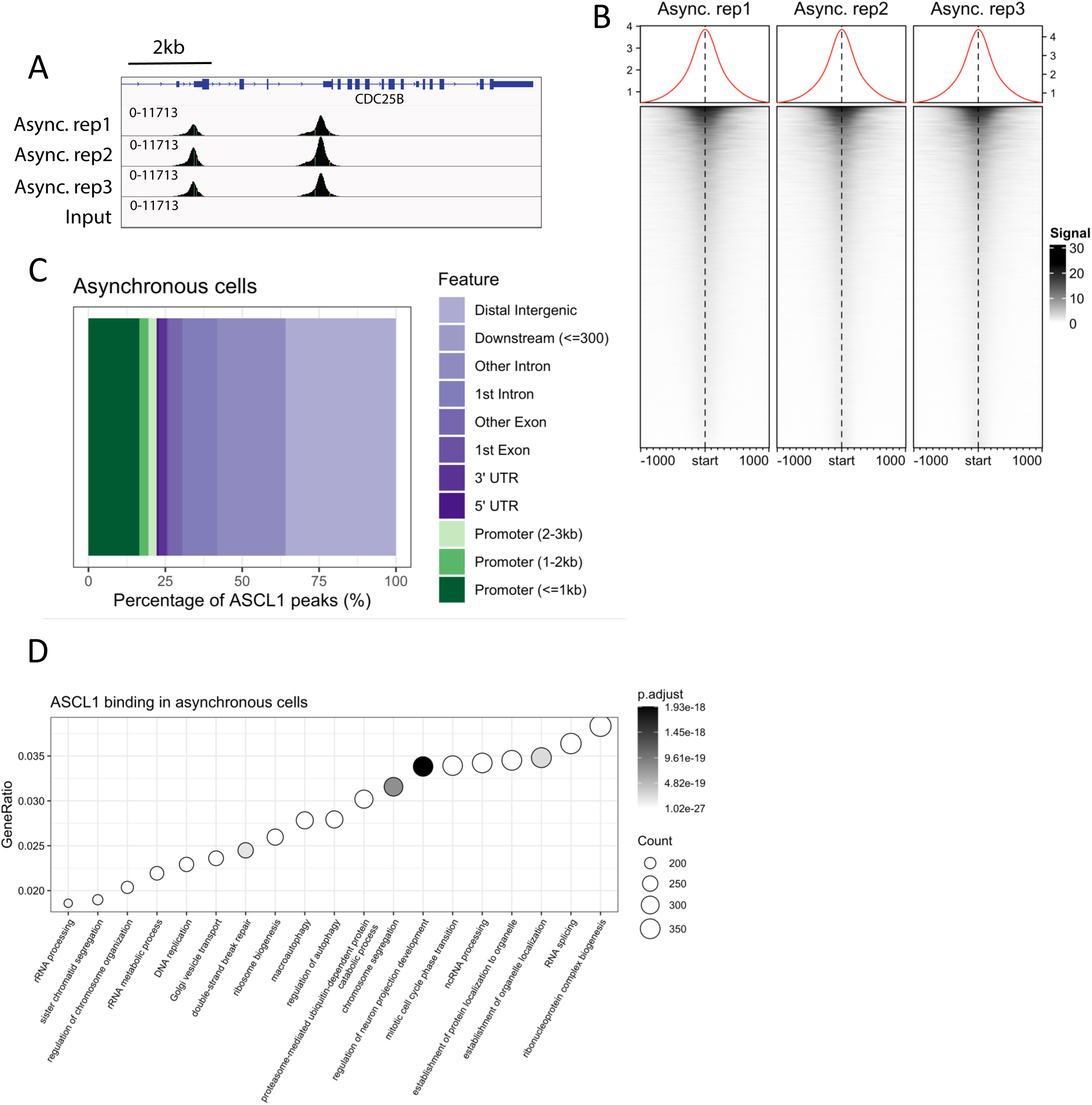
ASCL1 productively binds proliferative genes and non-productively binds neuronal genes. (A) ASCL1 ChIP-seq tracks of biological replicates in asynchronous SK-N-BE(2)-C cells, plus input control. (B) ASCL1 ChIP-seq heatmaps for three asynchronous replicates over the consensus peak set. (C) ASCL1 ChIP-seq consensus peak set annotation in asynchronous cells coloured by feature. (D) Gene ontology analysis of ASCL1 ChIP-seq peaks in asynchronous cells annotated to their nearest expressed gene TSS within 50kb.

To ascertain whether ASCL1 binding at these loci regulated gene expression in cycling neuroblastoma cells, we generated two ASCL1 knockout SK-N-BE(2)-C clones and validated ASCL1 protein absence by western blot (Figure 2A). While there were no obvious morphological changes in these cells (Figure S1C, D), knockout of ASCL1 significantly reduced the growth rate when compared to wild-type cells (Figure 2B), indicating that ASCL1 indeed plays a role in promoting cell cycle progression, as has been seen in other neuroblastoma cell lines^12^. We next performed RNA-seq on the parental cell line and one of the ASCL1 knockout clones (CRISPR 1), identifying 4474 genes that are significantly downregulated after ASCL1 knockout (Figure 2C). Gene ontology analysis of these downregulated genes (Figure 2D) uncovered similar terms to those identified from the ASCL1 ChIP-seq peak set including RNA and ribosomal terms, strongly supporting a role for ASCL1 in regulating these biological processes by directly binding constituent genes and their regulatory elements. Interestingly, none of the top 50 terms identified following gene ontology analysis of downregulated genes related to neuronal processes or functions, indicating that ASCL1 binding is not required for expression of neuronal genes in asynchronous freely cycling cells. However, many pro-proliferative and mitotic terms, including DNA replication and sister chromatid separation, were identified in the gene ontology analysis of downregulated genes, suggesting that ASCL1 actively drives cell cycle progression in proliferating neuroblastoma cells.

**Figure 2.**
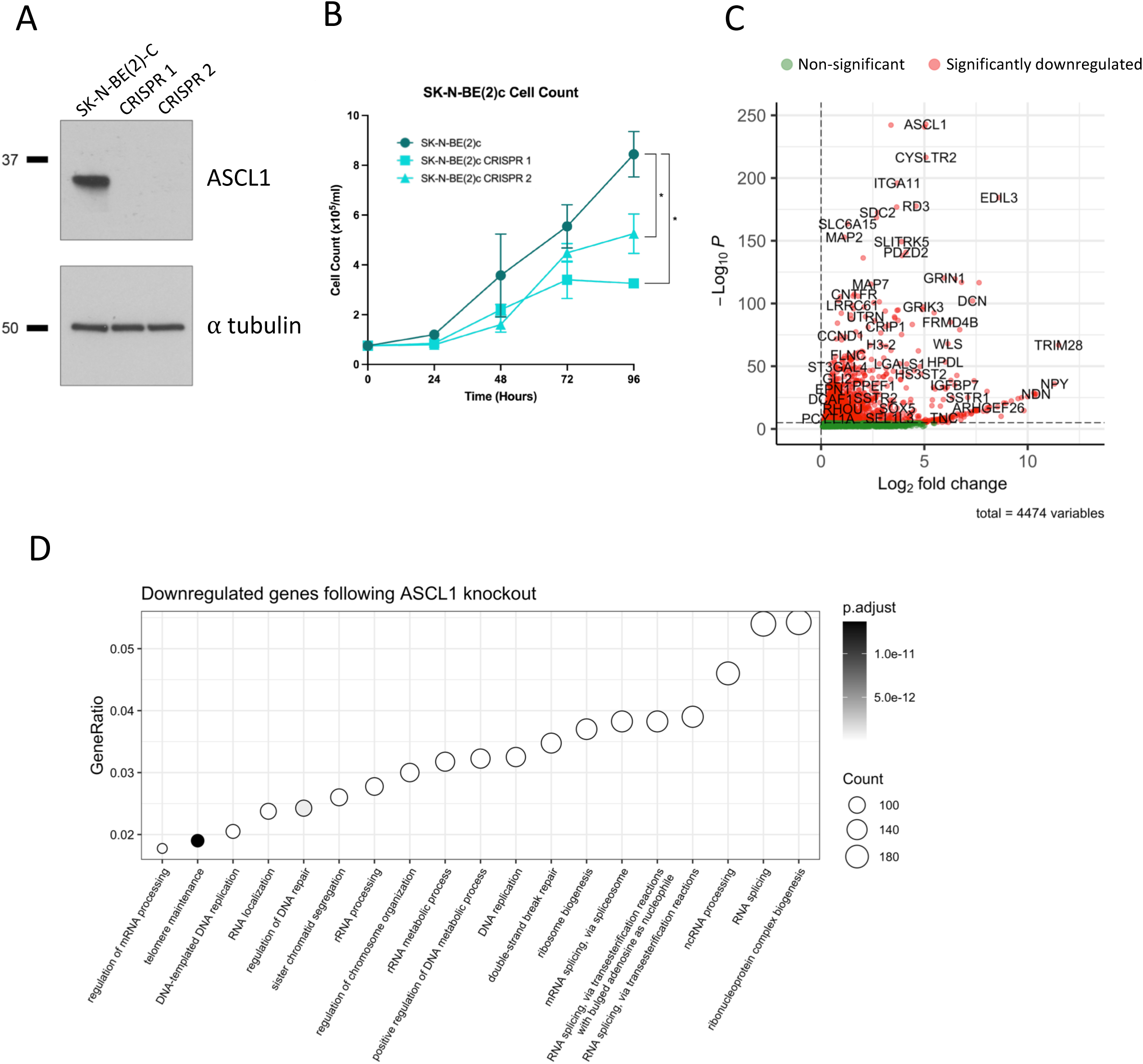
ASCL1. drives cell cycle progression in proliferating neuroblastoma cells (A) ASCL1 western blot in parental SK-N-BE(2)-C cells, and two ASCL1 CRISPR knockout clones. (B) Cell proliferation assay for SK-N-BE(2)-C parental cells and both ASCL1 CRISPR knockout clones. Mean values are shown (n=3) and SEM (unpaired two-tailed t-test; *=p<0.05). (C) RNA-seq results comparing ASCL1 WT SK-N-BE(2)-C and the ASCL1 knockout clone. Only genes showing a fold change of > 0 are shown, with significant genes in red and non-significant genes in green. (D) Gene ontology analysis of downregulated genes following ASCL1 CRISPR knockout.

### ASCL1 binding profiles in transiently synchronised cells recapitulates binding dynamics in cycling cells

Given that SK-N-BE(2)-C cells represent immature neuronal precursors and ASCL1 has been extensively investigated as a neurogenic factor ^38–40^, we sought to elucidate how ASCL1 can bind to and promote the expression of pro-proliferative genes while simultaneously binding neuronal loci without engaging a neuronal differentiation programme. We hypothesised that the binding profile determined from asynchronously cycling cells may represent two or more distinct ASCL1 binding profiles, each being derived from cells at different phases of the cell cycle. To test this, we generated a SK-N-BE(2)-C cell line stably harbouring the FUCCI cell cycle dual reporter system, allowing us to assign individual cells to distinct cell cycle phases, and monitor cell cycle progression ^41^. The FUCCI system consists of a green fluorophore (mAG) tagged with a geminin-derived degron, and a red fluorophore (mKO2) tagged with a cdt1-derived degron. Hence cells will acquire an overall red fluorescence when they are in (mid-late) G1 phase, and a green fluorescence during G2 and M phases. Cells in early G1 phase will accumulate neither the red nor the green fluorophore whereas cells cycling through S phase will exhibit both red and green fluorescence and appear yellow, while stalling cells in S phase will result in overall green fluorescence (Figure 3A) ^42,43^.

**Figure 3.**
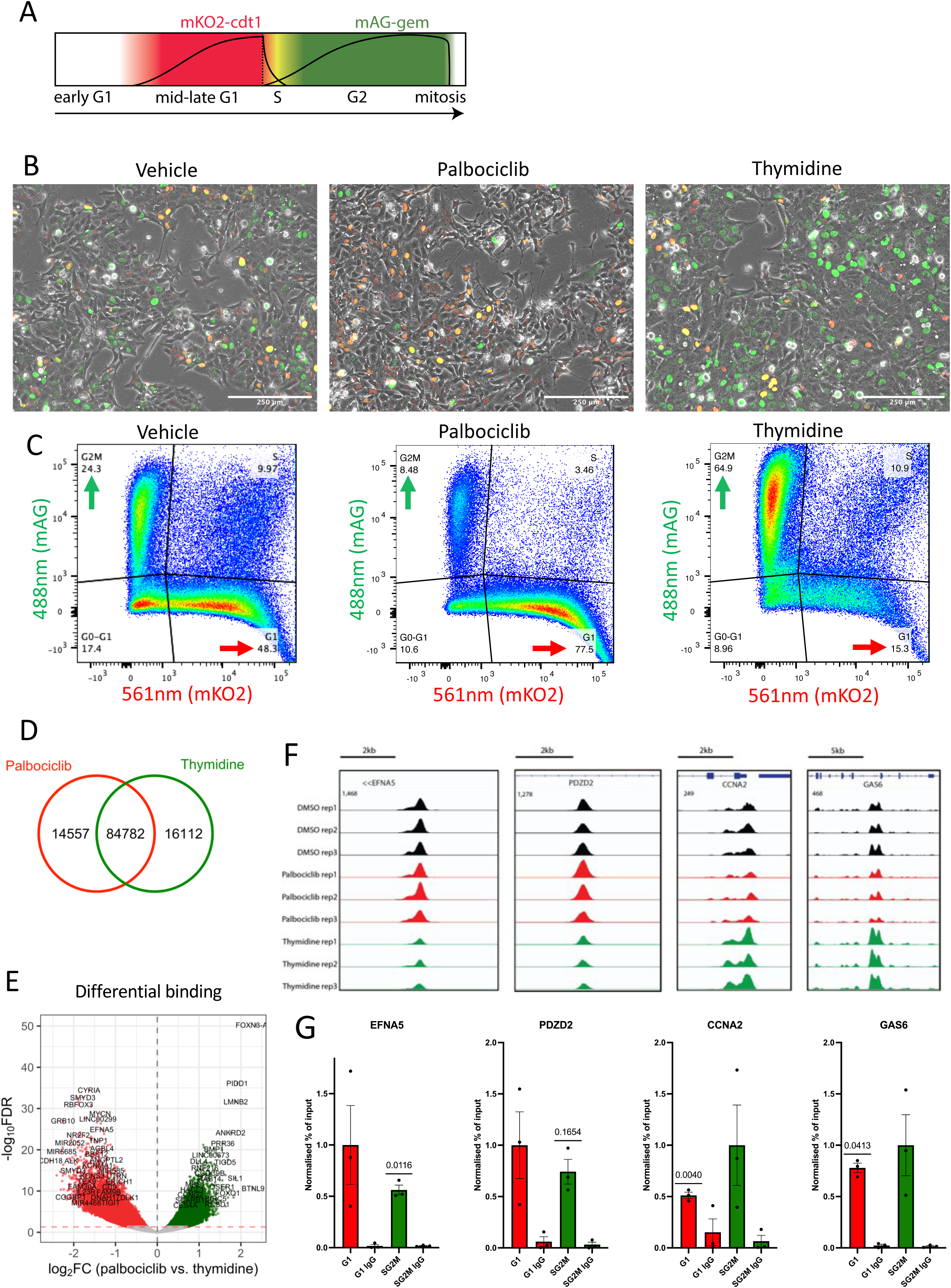
Cell cycle synchronisation mirrors ASCL1 binding dynamics in cycling cells. (A) Schematic diagram of FUCCI cell cycle reporter system. (B) Fluorescence images of SK-N-BE(2)-C FUCCI reporter cells following DMSO (left), palbociclib (centre) and thymidine (right) treatments. Scale bars represent 250 μm. (C) Flow cytometry analysis of SK-N-BE(2)-C FUCCI reporter cells after DMSO (left), palbociclib (centre) and thymidine (right) treatments. (D) Venn diagram depicting ASCL1 ChIP-seq consensus peaks overlap between palbociclib and thymidine treated cells. (E) Volcano plot of ASCL1 ChIP-seq peaks following DiffBind analysis. Red dotted line represents -log10(0.05) p-value. Green points represent thymidine enriched loci, and red points represent palbociclib enriched loci. (F) Replicate ASCL1 ChIP-seq tracks for DMSO (black), palbociclib (red) and thymidine (green) treated cells around two palbociclib (left) and two thymidine (right) enriched sites. (G) ASCL1 ChIP-qPCR for the four sites shown in (E) on FACS sorted G1 or SG2M cells. (see methods). Error bars show SEM (n=3, one sample two tailed t-test).

We used the CDK4/6 inhibitor, palbociclib, to stall the cells in G1 phase resulting in an increase in mKO2 positive cells, and thymidine to stall the cells in S phase resulting in an increase in mAG positive cells. (Figure 3B and 3C). DNA staining and karyotyping using Hoechst 33342 revealed that palbociclib treatment reduced the fraction of cells with replicated DNA (4c) characteristic of G2M phase cells, while thymidine treatment resulted in an accumulation of cells with a DNA content consistent with those expected in S phase cells, where chromosomes have been partially replicated (Figure S2A). Removal of both drugs resulted in detectable re-entry into the cell cycle within 3 hours (Figure S2B, S2C and S2D), demonstrating cell viability and drug reversibility ^44,45^.

After stalling cells in G1 or S phase, inhibitors were washed out to mitigate any direct effects of the drugs on transcription factor binding, and ChIP-seq for ASCL1 was performed on the enriched populations fixed 45 minutes post-drug removal. The majority of ASCL1 peaks were common between the palbociclib- and thymidine-stalled cells (Figure 3D), but DiffBind analysis identified 47,294 differentially bound sites between the two conditions (Figure 3E). Four of these sites (EFNA5 and PDZD2, enriched in palbociclib samples, and CCNA2 and GAS6, enriched in thymidine samples) were selected for validation by ChIP-qPCR in populations of cells that were FACS sorted according to cell cycle stage (Figure 3F), based on expression of the FUCCI reporters. Three out of 4 target sites showed significant cell cycle stage-specific binding that recapitulated the differential pattern seen after palbociclib or thymidine-induced stalling, while the fourth site showed a trend consistent with this differential binding, but did not meet statistical significance (Figure 3G). We therefore concluded that the cell cycle synchronisation approach using drugs reveals cell cycle-dependent differences in ASCL1 binding that are also seen at different cell cycle stages in asynchronously cycling cells.

### ASCL1 preferentially binds to neuronal loci during G1 phase, and pro-proliferative loci during SG2M phase of the cell cycle

We next sought to determine the potential significance of cell cycle stage-specific binding of ASCL1 to the genome. Of the 47,294 sites identified as differentially bound by ASCL1 in G1 versus SG2M cells, 32,834 sites showed higher levels of ASCL1 binding during G1 phase, and 14,460 showed higher levels of ASCL1 binding during SG2M. In asynchronous cells, peaks enriched for ASCL1 binding in SG2M showed generally higher levels of ASCL1 binding compared to those enriched in G1, which was replicated in two other neuroblastoma cell lines (Figure S3). We then defined all peaks with a false discovery rate (FDR) of over 0.05 as showing cell cycle-independent binding between the two cell cycle stages. All the peaks in these three subsets (G1 enriched, SG2M enriched and cell cycle-independent) were assigned to the nearest expressed gene TSS using a 50kb distance cut-off. These three gene sets were found to be non-mutually exclusive, as many genes associated with G1 or SG2M enriched peaks also had cell cycle-independent peaks (Figure 4A). Gene ontology analysis of associated genes revealed that that G1 enriched peaks were associated with genes involved in axonogenesis and neuronal differentiation processes, including small GTPase-mediated signal transduction and actin filament organisation, which are crucial for neuronal morphogenesis and polarity ^46,47^. In contrast, SG2M enriched peaks were overwhelmingly associated with pro-proliferative and mitotic genes rather than neuronal genes. Cell cycle-independent peaks were associated with more varied functions including metabolic and ribosomal, but also some neuronal processes (Figure 4B).

**Figure 4.**
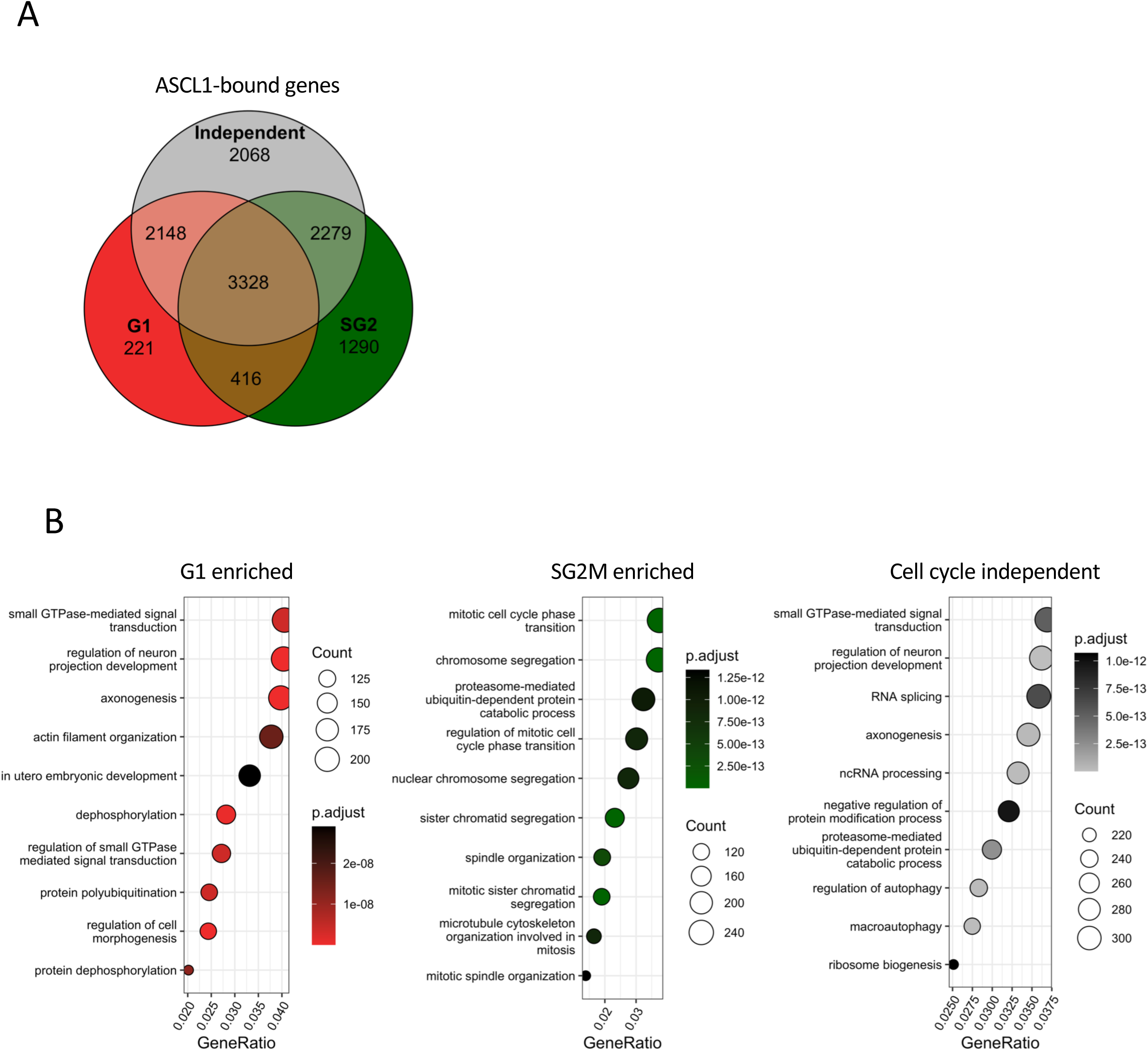
ASCL1 binds neuronal genes during G1 phase and proliferative genes during SG2M phase of the cell cycle. (A) Venn diagram of genes with G1 enriched (red), SG2M enriched (green) or cell cycle independent (grey) ASCL1 ChIP-seq associated peaks. (B) Gene ontology analysis of G1 enriched (red), SG2M enriched (green) and cell cycle independent (grey) ASCL1 ChIP-seq peaks. Only peaks within 50kb of an expressed gene were used.

To investigate the dependency of these genes on ASCL1 binding for expression, we took genes in the top 10 GO terms associated with ASCL1 binding in G1, SG2M or exhibiting cell cycle-independent binding, and analysed their expression in the ASCL1 knock-out clone compared to cycling parental cells expressing endogenous ASCL1. The gene sets associated with cell cycle-independent ASCL1 binding or those with enriched ASCL1 binding during SG2M were significantly downregulated in ASCL1 knockout cells, indicating that they normally rely on endogenous ASCL1 binding for expression. In contrast, genes with enriched ASCL1 binding during G1, including neuronal genes, did not show a significant decrease in expression level after ASCL1 knock-out (Figure 5A). This indicates that ASCL1 is capable of binding to neuronal targets in G1 phase of the cell cycle in neuroblastoma cells but is not supporting their expression under cycling conditions.

**Figure 5.**
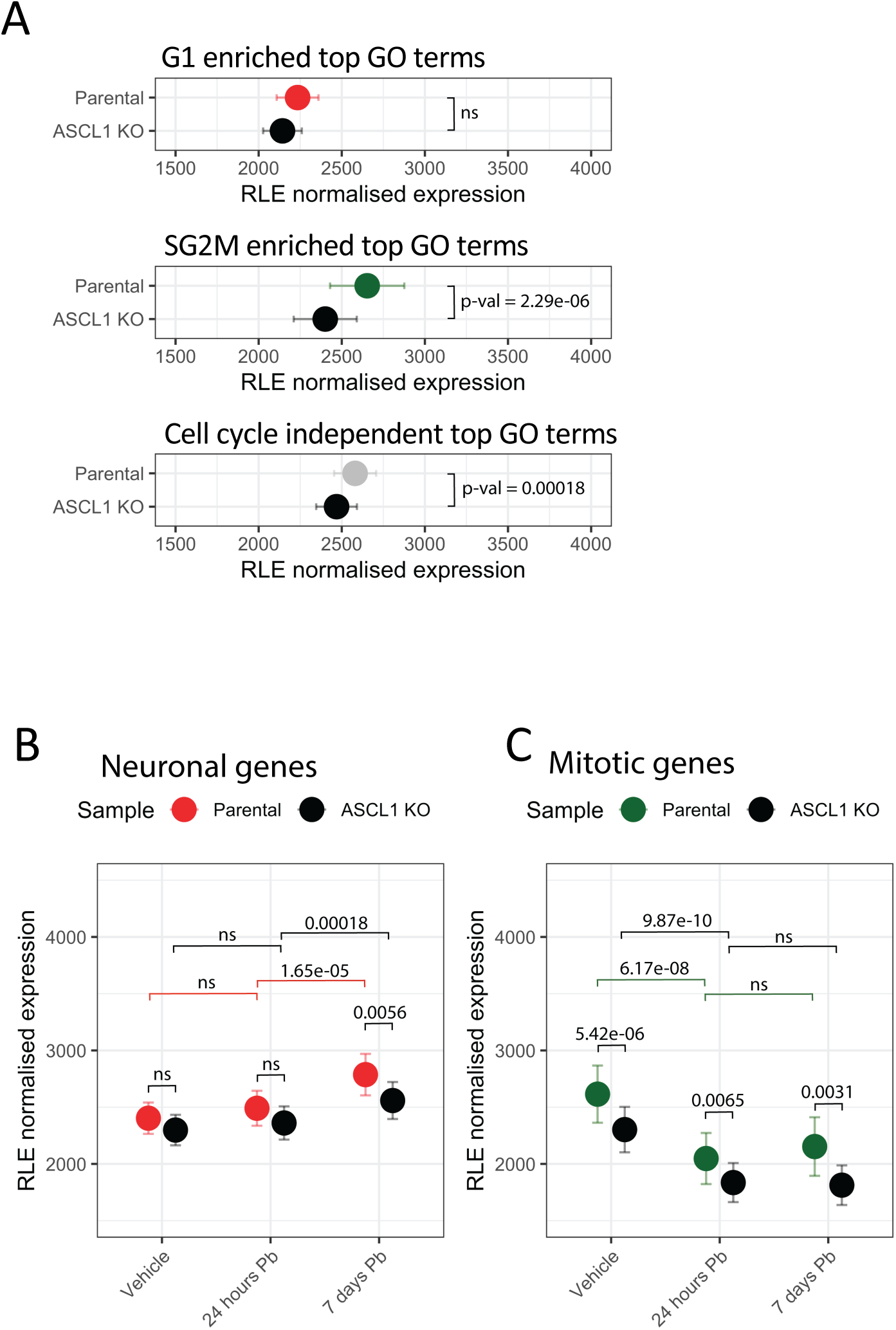
Ascl1 dependent switch from mitotic to neuronal specific gene expression in G1 arrested neuroblastoma cells. (A) RNA-seq expression of genes derived from the top 10 GO terms from Figure 4B in parental (coloured) or ASCL1 knockout (black) cells. Error bars represent SEM (merge of four replicates, two tailed t-test). (B) RNA-seq expression in parental (red) or ASCL1 knockout (black) cells for genes derived from neuronal GO terms of G1 enriched ASCL1 ChIP-seq peaks following palbociclib treatment. Error bars represent SEM (merge of four replicates, two tailed t-test). (C) RNA-seq expression in parental (green) or ASCL1 knockout (black) cells for genes derived from mitotic GO terms of SG2M enriched ASCL1 ChIP-seq peaks following palbociclib treatment. Error bars represent SEM (merge of four replicates, two tailed t-test).

### Prolonged G1 arrest converts primed ASCL1 binding at neuronal genes into productive binding

Differentiation is associated with an elongated G1 phase ^24^, and we have previously shown that G1 arrest can trigger a re-engagement of the differentiation process in neuroblastoma cells ^23^. Hence, we hypothesised that in G1 phase of cycling cells, ASCL1 may be marking neuronal genes for potential expression, but the short duration of G1 phase in these neuroblastic cells may be insufficient to allow ASCL1 binding to activate neuronal gene expression.

To investigate whether prolonged arrest in G1 phase is required to enable ASCL1 to activate expression of these ASCL1-bound, G1-enriched neuronal genes, we performed RNA-seq after 24 hours and 7 days of palbociclib treatment, comparing gene expression in ASCL1 WT and knock-out cells. Focussing on genes associated with neuronal functions that are preferentially bound by ASCL1 during G1 phase compared to SG2M, we found that their expression was significantly increased after 7 days of palbociclib treatment (Figure 5B). Crucially, expression of this gene set was not significantly different in ASCL1 knock-out cells growing asynchronously (Figure 3C, vehicle) or after 24 hours of palbociclib treatment. However, cells showed significantly more upregulation of neuronal genes after 7 days of palbociclib treatment when ASCL1 is present, indicating that ASCL1 cannot drive expression of these neuronal genes in neuroblasts unless G1 arrest is prolonged. We also looked at the pro-mitotic genes preferentially bound by ASCL1 during SG2M phase and found that, in contrast to neuronal genes, their expression was ASCL1-dependent in asynchronously cycling cells (Figure 3D, vehicle). As expected, their expression was significantly reduced by 24 hours and remained low after 7 days of palbociclib treatment (Figure 5C), reflecting the rapid exit of cells from the cell cycle following CDK inhibition. Our data reveal that ASCL1 binds neuronal genes specifically in G1 phase, but it does not drive expression of these genes in cycling neuroblasts, instead requiring prolonged G1 arrest for neuronal gene activation.

### ASCL1 preferentially interacts with inaccessible distal regulatory elements during G1 phase, and accessible promoter elements during SG2M phase of cycling cells

Given the different nature of the genes targeted by ASCL1 in G1 and SG2M and the differential reliance on endogenous ASCL1 in cycling cells to support their expression, we wanted to investigate the genomic context of ASCL1 enriched binding sites between G1 and SG2M. We returned to the previously identified cell cycle-independent, G1-enriched and SG2M-enriched ASCL1 ChIP-seq peaks (Figure 4B) and investigated peak locations in relation to known gene positions and TSSs. We found 15% of cell cycle-independent peaks were found within 3kb of a known TSS while 85% were located in exonic and intergenic regions (Figure 6A). Surprisingly, G1 enriched peaks were almost entirely annotated to non-promoter regions with only approximately 5% of binding sites occurring within 3kb of a known TSS, whereas the composition of SG2M peaks differed substantially, with approximately 60% of SG2M enriched ASCL1 binding occurring around a known TSS (Figure 4A).

**Figure 6.**
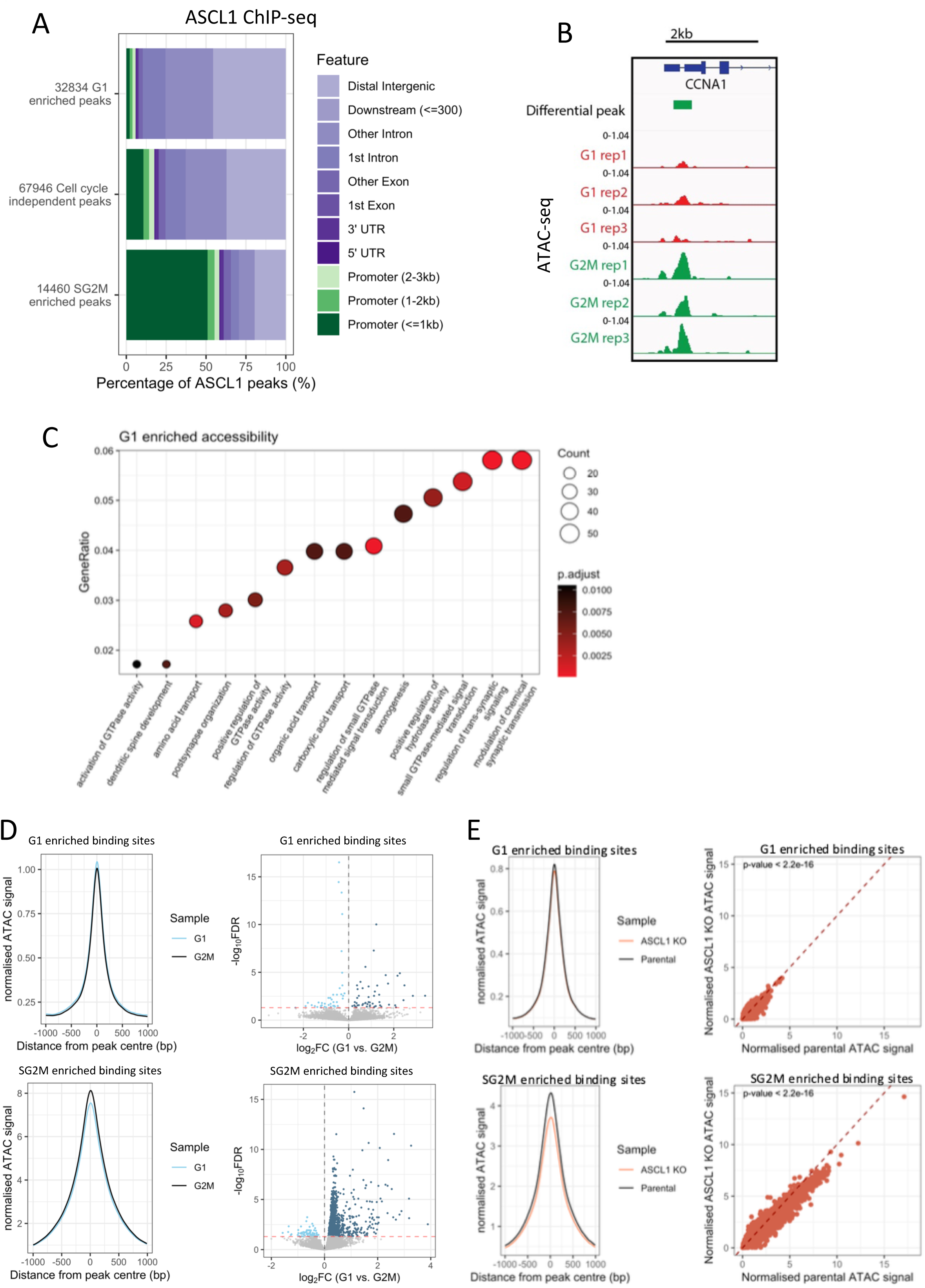
ASCL1 preferentially binds closed enhancers during G1 phase and open promoters during SG2M phase of the cell cycle. (A) ASCL1 ChIP-seq peak annotation for G1 enriched (top) cell cycle independent (middle) and SG2M enriched peaks (bottom). (B) Normalised ATAC-seq tracks for G1 (red) and G2M (green) replicates around CCNA1 promoter. (C) Gene ontology analysis of G1 enriched accessible regions following annotation to their nearest TSS within 50kb. (D) Normalised ATAC-seq signal for G1 (blue) and G2M (black) sorted cells at ASCL1 bound, G1 enriched (top panels) and SG2M enriched peaks (bottoms panels). Right panels show average ATAC-seq signals in G1 or G2M sorted cells for each differential ASCL1 ChIP-seq peak (two tailed, paired t-test). Dotted lines represent x=y. (E) Normalised ATAC-seq signal for parental SK-N-BE(2)-C (orange) and ASCL1 knockout (black) asynchronous cells at ASCL1 bound, G1 enriched (top panels) and SG2M enriched peaks (bottoms panels).

If ASCL1 can bind to distal regulatory elements during G1 phase but cannot activate associated genes in cycling cells, one might predict that accessibility levels at these sites would be low under these conditions, especially when compared to promoter regions of ASCL1-dependent genes bound preferentially by ASCL1 during SG2M. To test this, we FACS sorted cycling cells into enriched G1 phase or G2M phase populations based on DNA content and performed ATAC-seq (Figure 6B) to identify regions of the genome that show differential accessibility at different cell cycle phases. DiffBind analysis using a (FDR) corrected p-value threshold of 0.1 identified 1996 G1 phase and 1560 G2M phase enriched accessible sites (Figure S4A). Echoing our findings for differential ASCL1 binding sites at different cell cycle stages, we saw that sites that were more accessible during G1 phase were more likely to be distal from TSSs, while sites that were more accessible during G2M were more likely to be within 3kb of a TSS (Figure S4). Gene ontology analysis revealed G1 enriched accessible sites to be associated with genes involved in GTPase signalling and neuronal processes (Figure 6C), whereas G2M enriched accessible regions were not associated with any specific gene ontology category.

To ascertain whether ASCL1 binding affects differential accessibility at these sites at different cell cycle stages, we took all sites enriched for ASCL1 binding during G1, and all sites enriched for ASCL1 binding during SG2M and compared their accessibility profiles with ATAC-seq profiles from G1 and G2M cells. Chromatin regions at peaks showing higher levels of ASCL1 binding during G1 phase were more accessible during G1 phase than in G2M, while peaks showing higher levels of ASCL1 binding during SG2M phase were more accessible during G2M phase than during G1 phase. This indicates that ASCL1 binding does contribute to some modest differences in accessibility at different cell cycle stages. Nevertheless, on average, G1 ASCL1-bound sites had lower levels of accessibility than SG2M bound sites (Figure 6D). To validate a potential functional role of ASCL1 in contributing to accessibility at ASCL1-bound sites, we performed ATAC-seq on asynchronous ASCL1 WT or ASCL1 knockout cells. Analysis of the accessibility profiles at ASCL1-bound G1 phase enriched and SG2M phase enriched sites, revealed a significant decrease in accessibility at both the G1 sites and SG2M sites in the absence of ASCL1. However the change in accessibility at G1 sites was modest (Figure 6E) in agreement with the modest changes seen when comparing G1 and G2M accessibility (Figure 6D).

We have shown that ASCL1 binding in G1 is preferentially associated with genes associated with neuronal structures and functions (Figure 4B). This G1 binding is enriched at intergenic and intronic sites, many of which are likely to be potential enhancers, yet in cycling cells these are generally inaccessible (Figure 6D) and associated gene transcription is not maintained by ASCL1 (Figure 5B). Given that extended G1 cell cycle arrest can activate ASCL1-dependent expression of these genes, we hypothesised that the active epigenetic mark H3K27Ac would accumulate at these preferentially bound by ASCL1 sites during this extended G1 phase, supporting gene activation that does not occur in cycling cells. To test this, we analysed previously published H3K27ac ChIP-seq data in cycling SK-N-BE(2)-C cells ^23^ and compared accumulation of H3K27ac at sites with more ASCL1 binding in G1 compared to sites with more ASCL1 binding in SG2M (Figure 7A). There was an evident bimodal signal around the G1 enriched sites, which further supported their identity as potential enhancers ^48^, while the SG2M sites showed a less prominent bimodal peak, characteristic of active promoter elements ^48^. Next, we compared H3K27ac signal at cell cycle-enriched ASCL1 binding sites after 7 days of palbociclib treatment ^23^. G1 enriched ASCL1-binding sites showed significantly higher levels of H3K27ac after 7 days of palbociclib treatment compared to cycling cells and a more prominent bimodal distribution indicating an overall transition from ASCL1-bound inactive enhancers to active enhancers under these conditions. Conversely, SG2M enriched ASCL1-bound sites showed a significant reduction in H3K27ac after palbociclib treatment, mirroring the observed decrease in expression of these target genes (Figure 4) and likely deactivation of the pro-proliferative circuitry after G1 arrest. To test this hypothesis, we assessed the H3K27ac signal at the promoter elements of 2 canonical cell cycle genes before and after palbociclib treatment which indeed revealed a reduction in the mark at these sites following palbociclib treatment (Figure 7B), while the early neuronal marker, DCX, showed an increase in H3K27ac levels at the promoter and further downstream.

**Figure 7.**
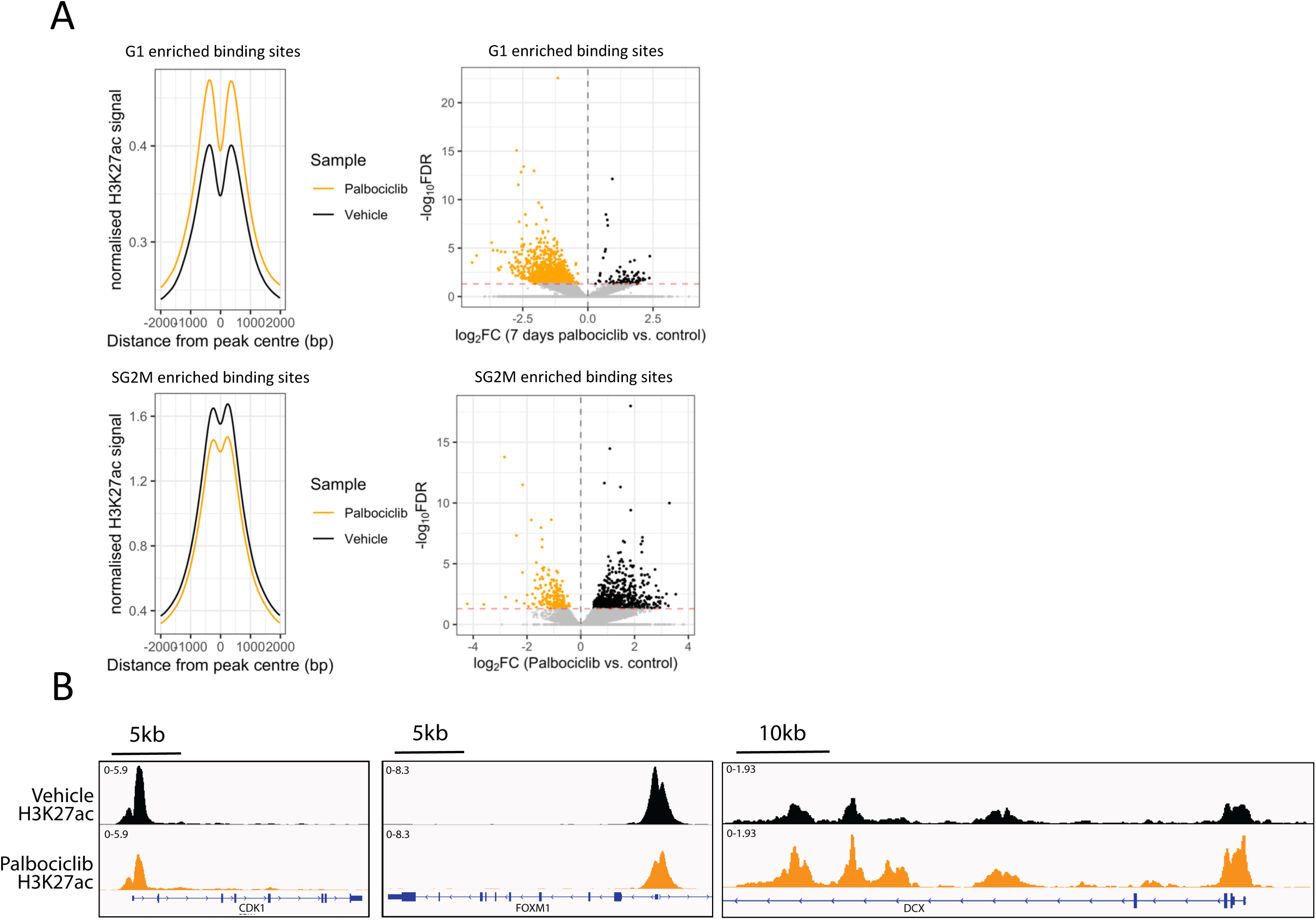
Prolonged G1 increases H3K27Ac mark at Ascl1 sites in Neuroblastoma cells. (A) Normalised H3K27ac ChIP-seq signal for control (black) and palbociclib treated (orange) SK-N-BE(2)-C cells at ASCL1 bound, G1 enriched (top panels) and SG2M enriched peaks (bottoms panels). Right panels show average H3K27ac ChIP-seq signal in control or palbociclib treated cells for each differential ASCL1 ChIP-seq peak (two tailed, paired t-test). Dotted lines represent x=y. (B) Merged (n=5) H3K27ac ChIP-seq tracks for control (black) and 7 days palbociclib treated (orange) cells around two cell cycle gene promoters (CDK1 and FOXM1) and one early neuronal gene (DCX).

## Discussion

During development, ASCL1 is a crucial yet enigmatic protein with well characterised roles in neuronal terminal differentiation and immature progenitor cell expansion. How one protein can direct these opposing biological processes has until now remained a mystery. In the present study we have addressed the hypothesis that in freely cycling cells, ASCL1 displays cell cycle phase dependent activity, revealing that G1 phase activity is associated more with neuronal differentiation, while SG2M activity associates more with progression of the cell cycle. Sites with preferential ASCL1 binding during G1 phase tend to be distally located enhancers of neuronal genes exhibiting low levels of accessibility and weak but bimodal H3K27ac peaks, indicative of primed enhancers ^27^. Sites with preferential binding during SG2M however tend to be promoter sites of pro-mitotic genes, exhibiting high levels of accessibility and more unimodal H3K27ac peaks. In these cycling cells, ASCL1 drives gene expression of the SG2M subset only, while prolonged G1 arrest is required to commission primed enhancers to drive associated neuronal gene expression.

How ASCL1 binding is regulated in a cell cycle phase dependent manner is yet to be elucidated, but we have previously shown that the phosphorylation of ASCL1 by CDKs is a key determinant of its differentiation potential ^34,36,49^ which itself could influence binding partners or degradation rate. CDK2 has previously been shown to phosphorylate and activate the histone methyltransferase, KMT2B, leading to H3K4me3 deposition selectively at developmental genes during late G1 phase in pluripotent stem cells ^50^. Co-occurrence of H3K4me3 and H3K27me3 defines developmental bivalent chromatin domains which are resolved during lineage specification such that associated genes are either activated, or permanently silenced ^51^. The significance of H3K4me3 at enhancer loci is still being debated ^51^, but one possibility here is that binding of ASCL1 at G1 enhancer sites is directed by G1 phase H3K4me3 deposition, and that prolonged G1 phase permits ASCL1 to resolve these bivalent domains.

That ASCL1 promotes pro-mitotic gene expression and cell cycle progression is at odds with our previous studies showing that ASCL1 overexpression drives potent neuronal differentiation in neuroblastoma cells ^35^. We propose that overexpression enables increased binding and aberrant activation of these ASCL1 bound G1 primed enhancers with corresponding target gene activation, while overexpression has little impact on the already activated cell cycle progression genes.

Taken together, our study reveals cell cycle stage-specific activities of ASCL1 and gives enhanced insight into how the length and structure of the cell cycle influences the balance between proliferation and differentiation of neuroblastic cells.

### Limitations of the study

Due to the required cell numbers for ChIP-seq, we used pharmacological cell cycle synchronisation methods to enrich cells in specific cell cycle stages. We made sure to remove the drugs with two PBS washes 45 minutes prior to performing ChIP-seq and validated the results in cycling asynchronous sorted cells, but there is always the possibility that the drugs had secondary effects on the activity of ASCL1. Additionally, we use a single neuroblastoma cell line as a model of ASCL1 activity, but this represents a pathological cellular state with many genetic and epigenetic aberrations. Thus it would be of interest to repeat this experiment in additional neuroblastoma lines and a more physiologically relevant context such as in neural stem cells.

## Acknowledgements

We would like to thank the NIHR Cambridge BRC Cell Phenotyping Hub for their support with FACS sorting; the Cambridge Stem Cell Institute Genomics and Tissue Culture facilities; Kirsty Ferguson, Lidiya Mykhaylechko, David Lando, and all of the Philpott lab for their useful discussions and feedback; Prof. Jason Carroll and Igor Chernukhin for their advice on ChIP-seq practices; Prof. Ludovic Vallier for providing FUCCI reporter plasmids.

A.P. was funded in whole, or in part, by the Wellcome Trust (203151/Z/16/Z, 203151/A/16/Z; 212253/Z/18/Z); Cancer Research UK (A25636) and the UKRI Medical Research Council (MC_PC_17230). For the purpose of open access, the author has applied a CC BY public copyright licence to any Author Accepted Manuscript version arising from this submission.

## Author contributions

Conceptualisation, W.B., A.P.; formal analysis, W.B.; writing – original draft, W.B.; writing – review & editing, W.B., A.P.; cloning, W.B.; ASCL1 CRISPR knockout, L.M.P.; ChIP-seq, W.B.; ATAC-seq, W.B., L.M.P.; RNA-seq, L.M.P., L.C.; imaging, W.B.; NGS data processing, W.B.; bioinformatic analysis, W.B.; visualisation, W.B.; supervision, A.P.; funding acquisition, A.P.; project administration, A.P.

---

**Supplementary Figure 1.**
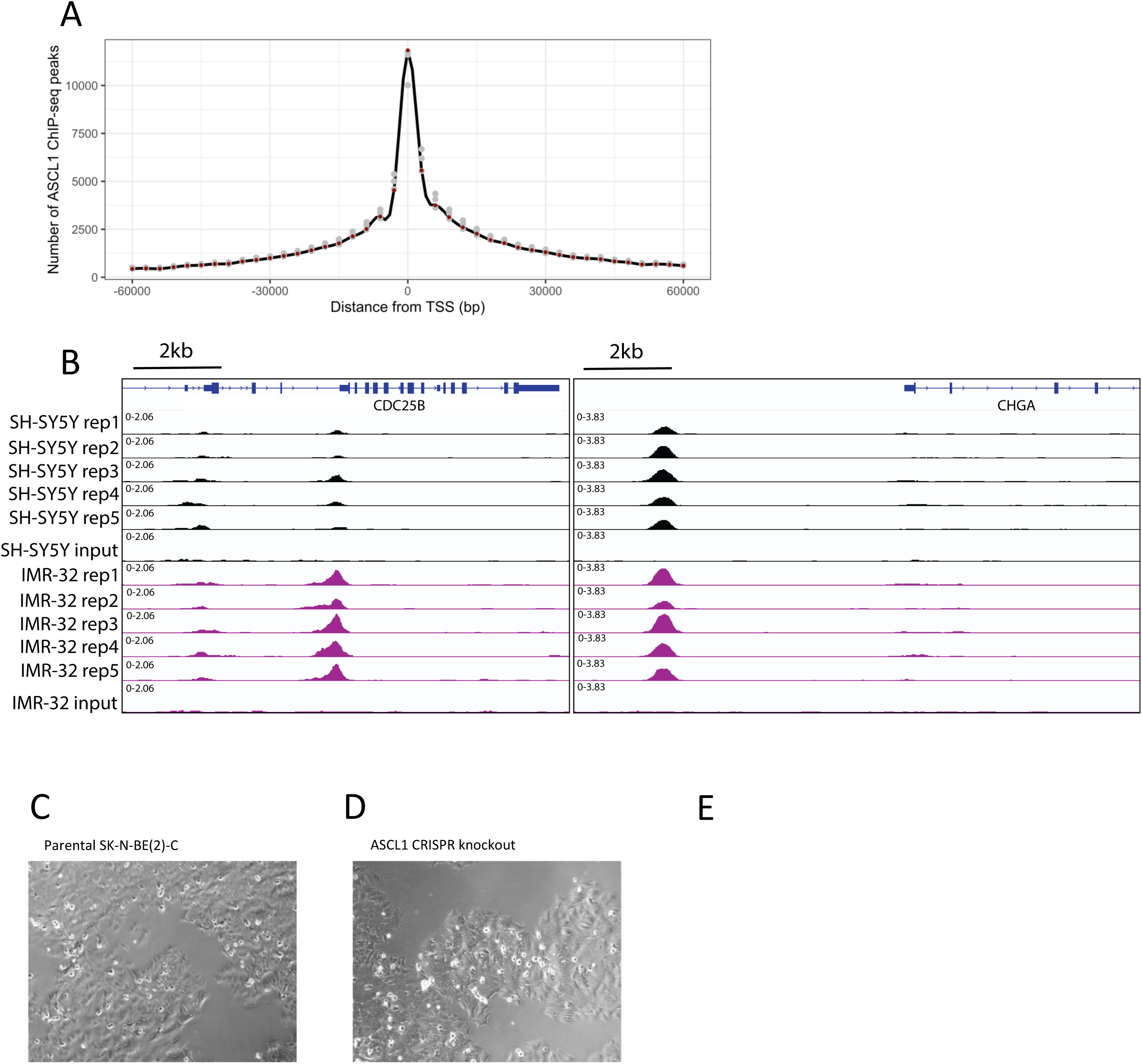
ASCL1 productively binds to pro-proliferative genes and non-productively binds to pro-neuronal genes. (A) Peak locations from 3 SK-N-BE(2)-C ASCL1 ChIP-seq replicates (grey) binned into 3kb windows from gene TSSs and quantified. Black line depicts the spline for the average across the replicates (red). (B) ASCL1 ChIP-seq tracks of five biological replicates in asynchronous SH-SY5Y cells (black) and IMR-32 cells (purple), plus the input controls. Regions around the CDC25B and CHGA TSSs are shown. (C) Phase contrast image of parental SK-N-BE(2)-C cell line. (D) Phase contrast image of the ASCL1 CRISPR knockout clone.

**Supplementary Figure 2.**
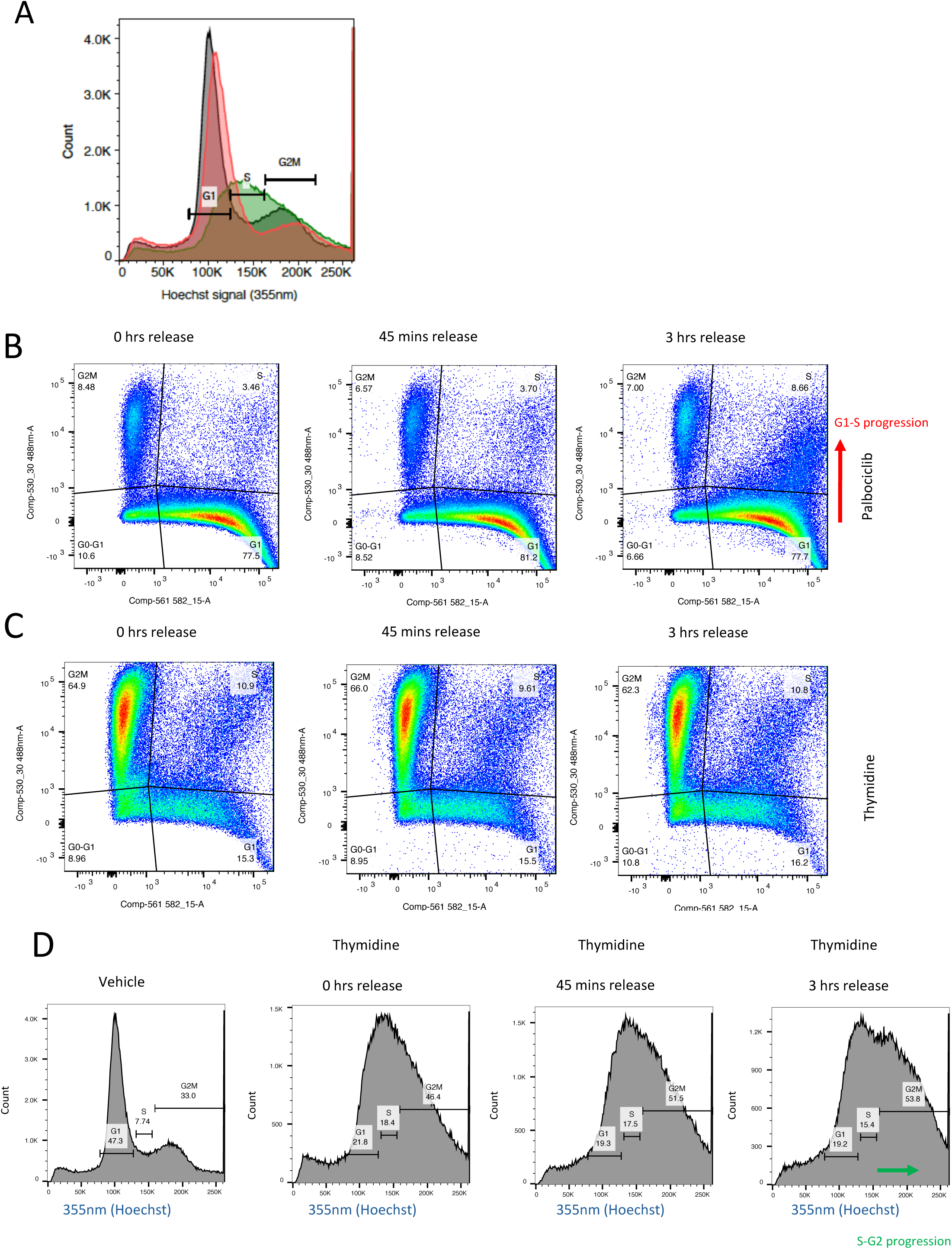
Cell cycle synchronisation is effective and reversible. (A) Flow cytometry plot of SK-N-BE(2)-C FUCCI cell line treated with DMSO (black), palbociclib (red) or thymidine (green), stained with Hoechst 33342. Cells are gated based on 355nm signal into G1 phase, S phase and G2M phase. (B) Flow cytometry and cell cycle analysis based on FUCCI fluorophores for cells treated with palbociclib followed by wash out for 0 mins (left), 45 mins (middle) and 3 hrs (right). X-axis shows mKO2 signal while y-axis shows mAG signal. (C) Flow cytometry and cell cycle analysis based on FUCCI fluorophores for cells treated with thymidine followed by wash out for 0 mins (left), 45 mins (middle) and 3 hrs (right). X-axis shows mKO2 signal while y-axis shows mAG signal. (D) Flow cytometry and cell cycle analysis based on DNA content for DMSO treated cells (left) and cells treated with thymidine followed by wash out for 0 mins (centre left), 45 mins (centre right) and 3 hrs (right). The same gates were used for all plots.

**Supplementary Figure 3.**
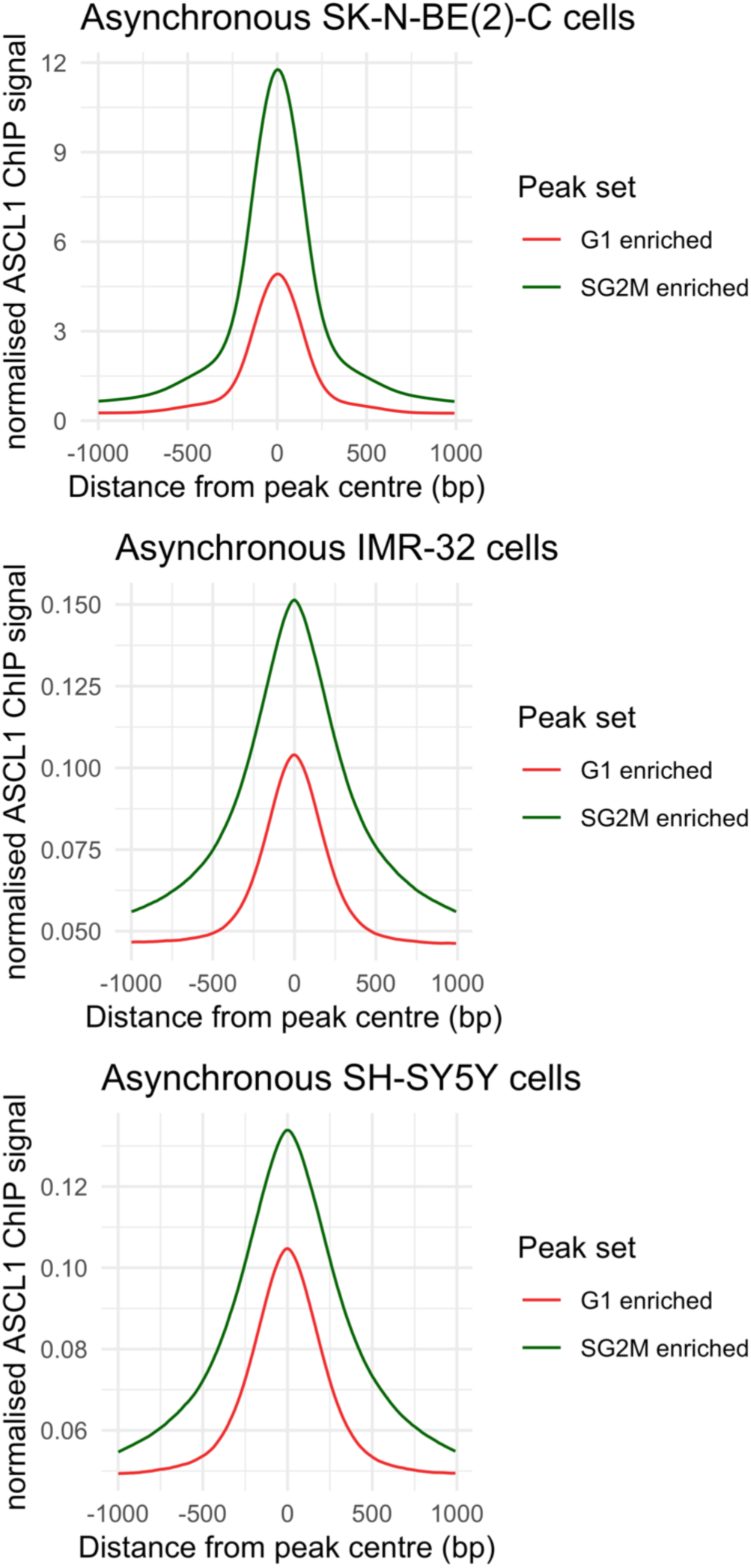
ASCL1 binding dynamics are consistent in additional neuroblastoma cell lines. Normalised ASCL1 ChIP-seq signal for peaks showing enriched ASCL1 binding in G1 phase (red) and SG2M phase (green) in asynchronous SK-N-BE(2)-C cells (top), IMR-32 cells (middle) and SH-SH5Y cells (bottom).

**Supplementary Figure 4.**
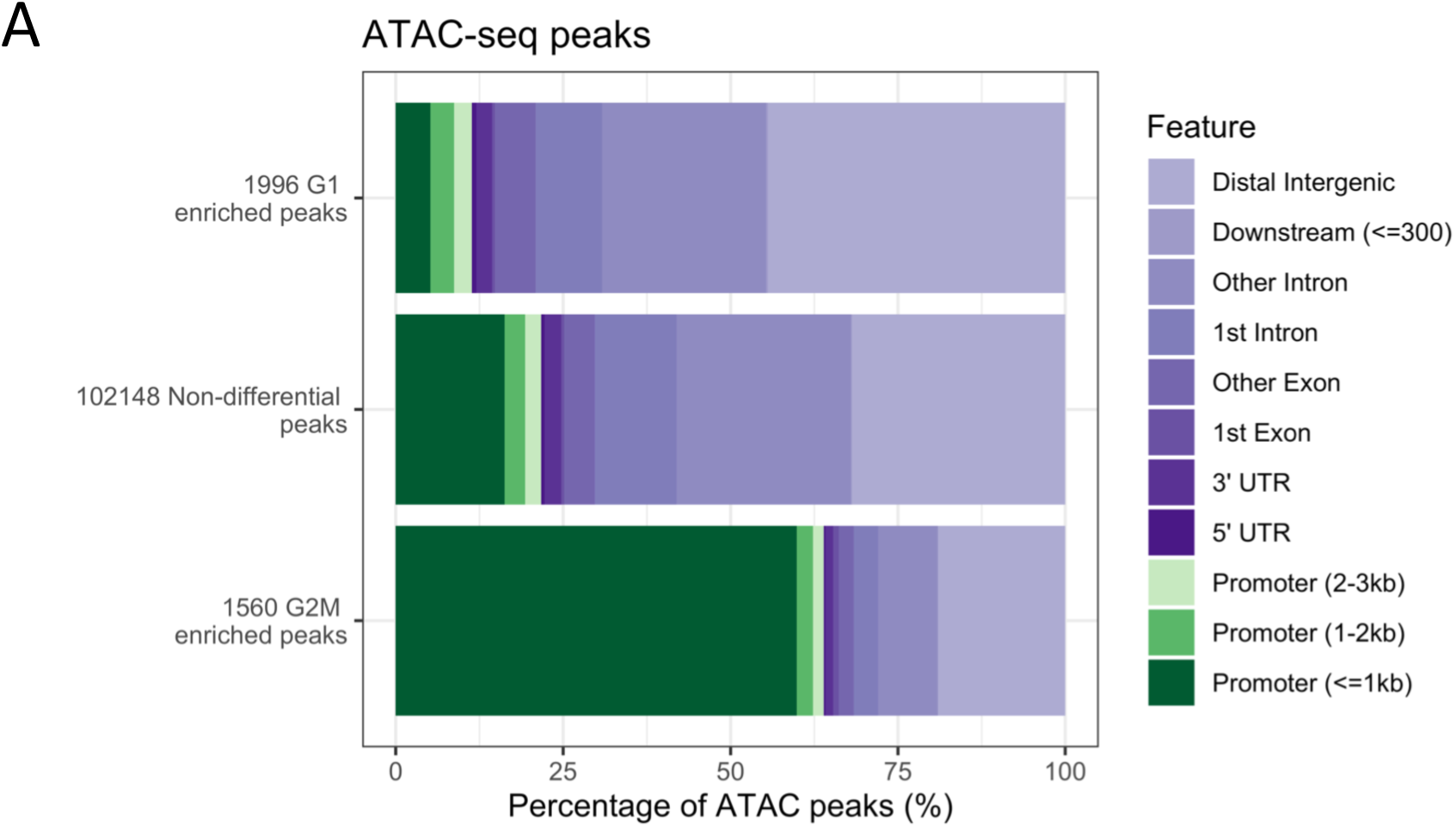
G1 enriched accessibility is associated with distal regulatory elements, while G2M enriched accessibility is associated with promoter elements. (A) G1 enriched (top), G2M enriched (bottom) and non-differential (middle) ATAC-seq peaks from FACS sorted populations. Peaks are coloured based on their genomic locations relative to known genes (green for promoter regions, purple for non-promoter regions).

## RESOURCE AVAILABILITY

### Lead contact

Further information and requests for resources and reagents should be directed to and will be fulfilled by the Lead Contact, Professor Anna Philpott.

### Materials availability

Plasmids generated in this study are available upon request.

### Data and code availability

ChIP-seq, ATAC-seq and RNA-seq data generated for this publication have been deposited at GEO and are publicly available as of the date of publication. Accession numbers are listed in the key resources table. All original code has been deposited at Zenodo and is publicly available as of the date of publication. DOIs are listed in the key resources table. Any additional information required to reanalyse the data reported in this paper is available from the lead contact upon request.

## EXPERIMENTAL MODEL AND STUDY PARTICIPANT DETAILS

### Cell culture

SK-N-BE(2)-C cells were cultured at 5% CO2 at 37°C in DMEM/F12 with L-glutamine and 15mM HEPES, and further supplemented with 10% FBS and 1% Penicillin-Streptomycin. Cells were tested regularly for mycoplasma infection by extended culture in the absence of Penicillin-Streptomycin followed by PCR. Cell lines were authenticated by short tandem repeat testing and comparison to the Cellosaurus database.

## METHOD DETAILS

### Generation of stable cell lines

Plasmids for the transient expression of the FUCCI dual reporter system were kindly gifted by Ludovic Vallier. GFP was removed from the 3rd generation pLenti-CMV-GFP-Hygro transfer plasmid using BamH1-HF and Sal1-HF and FUCCI reporters were amplified from transient expression plasmids for insertion using the Takara Bio In-Fusion cloning kit (see supplementary table for primer sequences). Lentiviral particles for the FUCCI reporters (pLenti-CMV-mAG-geminin, pLenti-CMV-mKO2-cdt1) were then generated in HEK293T cells. Briefly, 6µg of each plasmid harbouring TAT, GAG and POL, and REV, plus 12µg of VSV-G and 42µg of each transfer plasmid were added to a near confluent T175 flask of HEK293T cells in IMDM media. Resulting viral particles were collected, concentrated and quantified, and cells were transduced using a viral titre of 10 MOI in the presence of 8µg/ml polybrene. Single double positive cells were sorted from the transduced population and a stable clone was generated.

### Cell cycle synchronisation and ChIP-seq

3 million SK-N-BE(2)-C FUCCI reporter cells were plated in 10cm^2^ dishes then were either treated with DMSO for 24 hrs, 500nM palbociclib dissolved in DMSO for 24 hrs, or double blocked with 4mM thymidine for 24 hrs with a 12-hr release in between. After cell cycle synchronisation (or vehicle) all cells were washed twice with warm media then fresh media was added for 45 mins before cells were trypsinised and collected. Cells were pelleted and fixed in 1ml of cold solution A (1% formaldehyde; 50mM HEPES; 100mM NaCl; 1mM EDTA; 0.5mM EGTA) and fixation was stopped after 10 minutes then quenched with glycine to a final concentration of 125mM. Cells were washed twice with ice cold PBS supplemented with cOmplete protease inhibitor then pellets were frozen on dry ice and stored at -80°C.

50µl of protein G Dynabeads were used per samples and were washed twice in 1ml of 0.5 % BSA-PBS solution in protein LoBind tubes before being resuspended in 100µl 0.5 % BSA-PBS per sample. 4.5ug per sample of ASCL1 antibody was added. Bead-antibody solutions were then mixed and left to rotate for approx. 6 hours at 4°C.

ChIP-seq was performed as previously described ^53^. Briefly, cells were thawed then lysed in 1ml of cold lysis buffer 1 (50mM HEPES-KOH; 140mM NaCl; 1mM EDTA; 10% glycerol; 0.5% Igepal CA-630; 0.25% triton X-100) supplemented with cOmplete protease inhibitor then rotated at 4°C for 10 minutes before being centrifuged at 2000g for 5 minutes at 4°C. Pelleted nuclei were then resuspended in 1ml of cold lysis buffer 2 (10mM Tris-HCl; 200mM NaCl; 1mM EDTA; 0.5mM EGTA) supplemented with cOmplete protease inhibitor then rotated at 4°C for 10 minutes before again centrifuging at 2000g for 5 minutes at 4°C. Pelleted nuclei were then resuspended and lysed in 300µl of cold lysis buffer 3 (10mM Tris-HCl; 100mM NaCl; 1mM EDTA; 0.5mM EGTA; 0.1% Na-Deoxycholate; 0.5% N-lauroylsarcosine) supplemented with cOmplete protease inhibitor before being sonicated for 6 cycles in a Bioruptor 300 sonicator. 30µl of 10% Triton X-100 was then added to the lysate, and histone H2B and total mRNA added to final concentrations of 20ug/ml and 1ug/ml, respectively (acting as carrier). Lysate was centrifuged at 21000g for 10 minutes then supernatant was divided into 30µl for input, and 300µl for ASCL1 immunoprecipitation reaction. DNA LoBind tubes were used from this point on in the protocol. Bead-antibody mixes were removed from the cold room and washed 3 times with ice cold 0.5% BSA-PBS then resuspended in 100µl of 0.5% BSA-PBS per sample. 100µl of bead-antibody solution was added to the 300µl of lysate and mixed thoroughly. Both input and immunoprecipitation reaction were left to rotate overnight at 4°C.

The following day, immunoprecipitation samples were collected from the cold room and washed 10 times with cold RIPA buffer (50mM HEPES-KOH; 500mM LiCl; 1mM EDTA; 1% Igepal CA-630; 0.7% Na-deoxycholate) using a magnetic stand. On the last wash, samples were transferred to a fresh DNA LoBind tube and washed twice with cold TBS, before being resuspended in 200µl of elution buffer (50mM Tris-HCl; 10mM EDTA; 1% SDS). 170µl of elution buffer was added to the input samples and both were left at 65°C overnight on a shaker. The next day, supernatant was collected using a magnetic rack and all DNA samples were treated with RNaseA and proteinase K before being purified using phenol chloroform extraction.

### ChIP-seq sequencing and analysis

Replicate input samples were pooled at an equimolar ratio and sample libraries were prepared using the NEBNext Ultra II library prep kit. Libraries were then pooled and sequenced paired end to 150bp on a NovaSeq 6000. Adapter sequences were then trimmed using fastp ^54^ and reads mapped to the hg19 genome using Bowtie2 ^55^. Narrow peaks were called using MACS2 ^56^ with the DMSO input sample as the control and peaks with an enrichment score of less than 4 were removed. Differential peaks were identified using DESeq2 ^57^ in DiffBind ^58^ following reads in peaks library size normalisation using all peaks that were called in 2/3 replicates for each sample. ChIPseeker ^59^ was used to annotate peaks to their nearest TSS within 50kb unless otherwise stated. Gene ontology analysis was performed using clusterProfiler ^60^ using genes that showed an RLE normalised read count of >10 unless otherwise stated. Figures were made using ggplot2 ^61^ in R ^62^.

### ChIP-qPCR

SK-N-BE(2)-C FUCCI reporter cells were collected and fixed with cold solution A as described above. After quenching formaldehyde, cells were filtered and resuspended in FACS buffer (PBS with 5mM EDTA; 25mM HEPES: 0.1% BSA). 2 million green cells (SG2M) and 4 million non-green cells (G0/G1) were sorted into FACS buffer and ChIP was performed as described above, including an IgG control. Four differentially bounds sites identified from ChIP-seq were tested using primers designed to amplify small (approx. 90bp) regions. qPCRs were performed using PowerUp SYBR green master mix as per recommendations and run on an Applied Biosystems Stepone qPCR machine (see supplementary table for primer sequences).

### Generation of ASCL1 knockout line

This line was generated using the CRISPR-Cas9 system as previously described ^12^. The Cas9-2A-GFP and the U6-BsaI-sgRNA plasmids used for ASCL1 KO were kindly gifted by Prof. Steve Pollard (University of Edinburgh), the design and generation of these plasmids has been previously described ^63^ (see supplementary table for sgRNA sequence). Plasmids were transfected using Lipofectamine 2000 and cells were left to recover for 48 hrs. Clones were generated by FACS sorting single Cas9-GFP positive cells followed by expansion. Two clones were used, harbouring a 1 base pair deletion (c.883delC) and a 1 base pair insertion (c.793_794insG), or a 1 base pair deletion (c.883delC) and a 2 base pair insertion (c.882_883insGC). Absence of ASCL1 was confirmed by western blot.

### Western blot

Cells were collected by scraping into 1ml of ice-cold PBS and protein was extracted by resuspending cell pellet in RIPA buffer plus cOmplete protease inhibitor and incubating for 20 min on ice. Debris was removed by centrifuging at 16,000g for 10 min at 4°C and protein concentration was determined by BCA assay. An equal mass of protein was denatured in 1x NuPAGE LDS sample buffer with β-mercaptoethanol at 70°C for 10 minutes then loaded and separated on a BisTris gel with NuPAGE MOPS SDS running buffer at 150V. Protein was transferred to a nitrocellulose membrane over 1 hr at 4°C and 100V then the membrane was blocked for 1 hr using 5% milk. Primary antibodies were diluted in 1% milk and incubation took place overnight at 4°C. Membranes were washed thoroughly with TBST and secondary antibodies (anti-Mouse or anti-Rabbit HRP conjugated whole antibodies) were diluted in TBST then incubated with membranes at room temperature for 1 hr. Visualisation was performed using ECL western blotting detection reagent as per manufacturer’s instructions before being exposed to X-ray film.

### Proliferation assay

Cell lines were plated in duplicate, and cells were quantified on a CellCountess II Automated Cell Counter, using trypan blue cell stain to discriminate live and dead cell populations. Three biological replicates were measured per cell line.

### RNA-seq

ASCL1 WT SK-N-BE(2)-C and ASCL1 CRISPR knockout cells were plated at equal densities and treated with 1µM of palbociclib (or vehicle) for 24 hrs or 7 days. Cells were lysed *in situ* using RLT buffer and RNA was purified using the RNeasy mini kit (Qiagen) as per manufacturer’s instructions. Poly-A selection was performed, and libraries were made using NEBNext Ultra II directional RNA kit. Libraries for 5 biological replicates were pooled and 100bp paired end reads were generated on a NovaSeq 6000 (Illumina). Reads were trimmed and quality filtered using TrimGalore with a minimum phred score of 20 then aligned to hg19 using STAR ^64^ with quantMode to obtain read counts. DESeq2^57^ was used to normalise read counts and identify differentially expressed genes.

### ATAC-seq

For ATAC-seq of cell cycle FACS sorted cells, asynchronous SK-N-BE(2)-C we plated in 15cm^2^ dishes and grown to near confluency. Live cells were then stained with 3ug/ml Hoechst 33342 for 1 hour, before being trypsinised and collected. Cells were centrifuged for 5 minutes at 600g before being resuspended in FACS buffer (PBS with 5mM EDTA; 25mM HEPES: 0.1% BSA) and filtered with a 45µm filter. 50,000 G1 or G2M cells were sorted into 500µl FACS buffer using a FACSariaIII based on UV 355 signal. For ATAC-seq of asynchronous parental SK-N-BE(2)-C cells and ASCL1 knockout cells, 50,000 live cells from the total population were sorted using a FACSariaIII. For all samples, ATAC-seq was performed using the Omni-ATAC protocol as previously described ^65^. Briefly, cells were pelleted and resuspended in 1ml of cold resuspension buffer (RSB; 10mM Tris-HCl pH 7.4, 10mM NaCl, 3mM MgCl_2_). Cells were spun at 500g for 5 min at 4°C, then resuspended in 50µl of RSB containing 0.1% Tween-20, 0.1% Igepal CA-630 and 0.01% digitonin and incubated on ice for 3 min. 1ml of RSB containing only 0.1% Tween-20 was added and nuclei were centrifuged for 10 min at 500g at 4°C. Nuclei were then resuspended in 50µl of transposition mix (25µl 2x TD buffer, 2.5µl transposase, 16.5µl PBS, 0.5µl 1% digitonin, 0.5µl 10% Tween-20 and 5µl H_2_O) and incubated at 37°C for 30 min. DNA was then cleaned and purified using the Zymo DNA Clean and Concentrator-5 kit before being amplified and quantified on a Agilent tapestation. Three biological replicates of cell cycle sorted cells were prepared, pooled and sequenced on a NovaSeq 6000 to generate 100bp paired end reads. Four replicates of the asynchronous parental and ASCL1 knockout cells were prepared, pooled and sequenced on a NovaSeq 6000 to generate 50bp paired end reads. Resulting reads were trimmed using TrimGalore with a Phred score cut-off of 20 and then mapped to hg19 using Bowtie2 ^55^. MACS2 ^56^ was used to call peaks and DiffBind ^58^ (EdgeR^66^) for identifying differential peaks with reads in peaks library size normalisation. Bigwigs were scaled according to these scaling factors.

### Imaging

Fluorescent imaging was performed on an Olympus IX51 inverted microscope, while phase contrast only images were taken using an EVOS M5000.

## QUANTIFICATION AND STATISTICAL ANALYSIS

Statistical analyses were performed using Prism or R software^62^, with each statistical test, sample sizes and significance thresholds noted in the figure legends. Biological replicates were derived from cells at different passage numbers. Statistical analysis of differential binding and differential accessibility was performed using DiffBind^58^ in R and statistical analyses of RNA-seq data were performed using DESeq2^57^ in R ^62^.

## KEY RESOURCES TABLE

| REAGENT or RESOURCE | SOURCE | IDENTIFIER |
| --- | --- | --- |
| **Antibodies** |  |  |
| ASCL1 (ChIP-seq) | Abcam | ab74065 |
| ASCL1 (WB) | Abcam | ab211327 |
| IgG (ChIP-qPCR) | Abcam | ab171870 |
| α-Tubulin (WB) | ProteinTech | 66031 |
| Secondary antibody: Sheep anti-mouse IgG HRP conjugated whole Ab | GE Healthcare | NA931 |
| Secondary antibody: Donkey anti-rabbit IgG HRP conjugated whole Ab | GE Healthcare | NA934 |
| **Bacterial and virus strains** |  |  |
| Stellar competent cells | Takara Bio | 636763 |
| **Biological samples** |  |  |
| **Chemicals, peptides, and recombinant proteins** |  |  |
| DMEM/F12 | Sigma-Aldrich | D8437 |
| IMDM | Gibco | 21980032 |
| FBS | Pan BioTech | P40-37500 |
| Pen/Strep | Sigma-Aldrich | P0781 |
| Trypsin EDTA 0.25% | Gibco | 25200072 |
| DMSO | Santa Cruz Biotechnology | sc-358801 |
| BamH1-HF | New England Biolabs | R3136S |
| Sal1-HF | New England Biolabs | R3138S |
| Agarose | Sigma-Aldrich | A9539 |
| Gel red 1000x stain in DMSO | Biotium | 41002 |
| 6x purple loading dye | New England Biolabs | B7024S |
| DNA ladder | New England Biolabs | N0550S |
| Polybrene | Sigma-Aldrich | TR-1003-G |
| Palbociclib | Sigma-Aldrich | PZ0383 |
| Thymidine | Sigma-Aldrich | T9250 |
| Formaldehyde | VWR chemicals | 20909.290 |
| HEPES | Sigma-Aldrich | H3375 |
| NaCl | ThermoFisher Scientific | BP358-1 |
| EDTA | Millipore | 324503 |
| EGTA | Millipore | 324626 |
| Glycine | ThermoFisher Scientific | G/0800/60 |
| cOmplete protease inhibitor | Roche | 11836170001 |
| Dynabeads Protein G | Invitrogen | 10003D |
| BSA | Fisher BioReagents | BP9706-100 |
| KOH | Sigma-Aldrich | 757551 |
| Glycerol | ThermoFisher Scientific | G/0600/08 |
| Igepal CA-630 | MP Bio | 198596 |
| Triton X-100 | Sigma-Aldrich | X100 |
| Trizma base | Sigma-Aldrich | T1503 |
| HCl | Honeywell | 72033 |
| Na-Deoxycholate | Sigma-Aldrich | D6750 |
| N-lauroylsarcosine | Sigma-Aldrich | L5777 |
| Histone H2B | New England Biolabs | M2505S |
| Total RNA | ThermoFisher Scientific | 4307281 |
| LiCl | ThermoFisher Scientific | L/2201/50 |
| Proteinase K | Invitrogen | EO0491 |
| Phenol:chloroform:isoamylalcohol | Sigma-Aldrich | P3803 |
| Ethanol | Sigma-Aldrich | 32221-2.5L-M |
| RIPA buffer | Sigma-Aldrich | R0278 |
| BisTris gel | Invitrogen | NP0301 |
| Skimmed milk powder | Serva | 42590 |
| Tween-20 | Promega Corporation | H5151 |
| Nitrocellulose membrane | Bio-Rad | 1620115 |
| NuPAGE MOPS SDS running buffer | Invitrogen | NP0001 |
| NuPAGE LDS buffer | Invitrogen | NP0007 |
| β-mercaptoethanol | BDH | 441433A |
| Hoechst 33342 | Invitrogen | H3570 |
| MgCl<sub>2</sub> | Sigma-Aldrich | M1028 |
| Digitonin | Sigma-Aldrich | D141 |
| PowerUp™ SYBR™ Green Master Mix | Applied Biosystems | A25742 |
| Methanol | ThermoFisher Scientific | M/4056/15 |
| **Critical commercial assays** |  |  |
| ECL Western Blotting Detection Reagents | Cytivia | RPN2106 |
| CellCountess II Automated Cell Counters | ThermoFisher Scientific | C10312 |
| RNeasy mini kit | QIAGEN | 74104 |
| NEBNext Ultra II directional RNA kit | New England Biolabs | E7760L |
| Lipofectamine 2000 | ThermoFisher Scientific | 11668019 |
| Tagment DNA enzyme and buffer kit | Illumina | 20034198 |
| DNA Clean and Concentrator-5 kit | ZYMO | D4014 |
| D1000 Tapestation reagents | Agilent | 5067 |
| QuantiTect reverse transcription kit | Qiagen | 205311 |
| BCA protein assay kit | ThermoFisher Scientific | 23227 |
| Plasmid Maxi kit | QIAGEN | 12162 |
| **Deposited data** |  |  |
| ASCL1 ChIP-seq; raw and analysed data | This manuscript | GSE276842 |
| Cell cycle sorted ATAC-seq; raw and analysed data | This manuscript | GSE276994 |
| ASCL1 KO ATAC-seq; raw and analysed data | This manuscript | GSE276994 |
| RNA-seq; analysed data | This manuscript | GSE276843 |
| Neuroblastoma survival data | R2 Genomics Platform | Tumour Neuroblastoma public - Versteeg - 88 MAS5.0 - u133p2 |
| **Experimental models: Cell lines** |  |  |
| SK-N-BE(2)-C cells | Laboratory of Prof. Deborah Tweedle, Newcastle University | N/A |
| HEK293T cells | Cambridge Stem Cell Institute Tissue Culture facility | N/A |
| IMR-32 cells | Laboratory of Prof. Deborah Tweedle, Newcastle University | N/A |
| SH-SY5Y cells | Laboratory of Prof. John Hardy, UCL | N/A |
| **Experimental models: Organisms/strains** |  |  |
| **Oligonucleotides** |  |  |
| ChIP-qPCR primer sequences | Sigma-Aldrich | N/A |
| ASCL1 CRISPR sgRNA sequences | Laboratory of Prof. Steve Pollard, University of Edinburgh | N/A |
| In-Fusion primer sequences for Fucci reporter cloning | Sigma-Aldrich | N/A |
| **Recombinant DNA** |  |  |
| pCAG-mKO2-cdt1 | Laboratory of Prof. Ludovic Vallier | N/A |
| pCAG-mAG-gem | Laboratory of Prof. Ludovic Vallier | N/A |
| pLenti-CMV-mAG-gem | This study | N/A |
| pLenti-CMV-mKO2-cdt1 | This study | N/A |
| **Software and algorithms** |  |  |
| FIJI/Image J | Open Source | https://imagej.net/software/fiji/ |
| GraphPad Prism version 10.3.0 | GraphPad | https://www.graphpad.com/scientific-software/prism/ |
| StepOne Software | ThermoFisher Scientific | https://www.thermofisher.com/uk/en/home/technical-resources/software-downloads/StepOne-and- |
|  |  | StepOnePlus-Real-Time-PCR-System.html |
| TapeStation Software | Agilent | https://www.agilent.com/en/product/automated-electrophoresis/tapestation-systems/tapestation-software |
| IGV v2.17.4 | Robinson et al. <sup>67</sup> | https://igv.org/ |
| R Studio v 2024.04.0+735 | CRAN <sup>62</sup> | https://cran.r-project.org/ |
| ChIPpeakAnno v3.38.0 | Zhu et al. <sup>68</sup> | https://bioconductor.org/packages/release/bioc/html/ChIPpeakAnno.html |
| edgeR v4.2.0 | Robinson et al. <sup>66</sup> | https://bioconductor.org/packages/release/bioc/html/edgeR.html |
| Rtracklayer v1.64.0 | Lawrence et al. <sup>69</sup> | https://www.bioconductor.org/packages/release/bioc/html/rtracklayer.html |
| DiffBind v3.14.0 | Stark et al. <sup>58</sup> | https://bioconductor.org/packages/release/bioc/html/DiffBind.html |
| ChIPseeker v1.40.0 | Yu et al. <sup>59</sup> | https://github.com/YuLab-SMU/ChIPseeker |
| TxDb.Hsapiens.UCSC.hg19.knownGene v3.2.2 | Bioconductor | https://bioconductor.org/packages/release/data/annotation/html/TxDb.Hsapiens.UCSC.hg19.knownGene.html |
| DESeq2 v1.44.0 | Love et al. <sup>57</sup> | https://bioconductor.org/packages/release/bioc/html/DESeq2.html |
| clusterProfiler v4.12.0 | Yu et al. <sup>60</sup> | https://github.com/YuLab-SMU/clusterProfiler |
| org.Hs.eg.db v3.19.1 | Bioconductor | https://bioconductor.org/packages/release/data/annotation/html/org.Hs.eg.db.html |
| Enrichplot v1.24.0 | Bioconductor | https://www.bioconductor.org/packages/release/bioc/html/enrichplot.html |
| ggpubr v0.6.0 | CRAN | https://cran.r-project.org/web/packages/ggpubr/index.html |
| ggplot2 v3.5.1 | Wickham et al. <sup>61</sup> | https://cran.r-project.org/web/packages/ggplot2/index.html |
| Tidyverse v2.0.0 | CRAN | https://cran.r-project.org/web/packages/tidyverse/index.html |
| RColorBrewer v1.1-3 | CRAN | https://cran.r-project.org/web/packages/RColorBrewer/index.html |
| EnrichedHeatmap v1.34.0 | Gu et al. <sup>70</sup> | https://bioconductor.org/packages/release/bioc/html/EnrichedHeatmap.html |
| TrimGalore v0.6.10 | Babraham Bioinformatics | https://github.com/FelixKrueger/TrimGalore |
| STAR v2.6.1a | Dobin et al. <sup>64</sup> | https://github.com/alexdobin/STAR |
| Bowtie2 v2.5.4 | Langmead et al. | https://bowtie-bio.sourceforge.net/bowtie2/manual.shtml |
| MACS2 v2.2.7.1 | Zhang et al. | https://github.com/macs3-project/MACS |
| fastp v0.23.4 | Chen et al. | https://github.com/OpenGene/fastp |
| **Other** |  |  |
| X-ray film | Scientific Laboratory Supplies | MOL7016 |

